# A2AR-mediated inhibition of B cell maturation: a novel mechanism of immune suppression in cancer

**DOI:** 10.1101/2025.08.13.669666

**Authors:** Paola Tieppo, Hussein Shehade, Chiara Martinoli, Marjorie Mercier, David Carbonez, Nicolas Rosewick, Sophie Jung, Anais Vezzu, Boris Pirlot, Noemie Wald, Stephanie Ma, Lucas Chaible, Reece Marillier, Francesco Strozzi, Joao Marchante, Margreet Brouwer, Erica Houthuys, Michael Deligny, Yvonne McGrath, Maura Rossetti

**Affiliations:** iTeos Therapeutics, Gosselies, Belgium

**Keywords:** adenosine, A2AR, B cells, plasma cells, cancer

## Abstract

High levels of extracellular adenosine, highly abundant in the tumor microenvironment, promote immune suppression mainly through the A2AR expressed by tumor-infiltrating immune cells. Here we show that plasma cells are the most negatively affected by adenosine among the immune cells present in the tumor microenvironment. Furthermore, both tonsillar and tumor-associated B cells, including germinal center (GC)-like B cells, plasma cells and plasma blasts (PCs and PBs, respectively, and collectively referred to as antibody-secreting cells (ASCs), express high levels of A2AR. Given the importance of tumor-infiltrating B and PCs in antitumor responses, we investigated how adenosine impairs their numbers and function. Triggering of A2AR inhibited B cell maturation into ASCs and immunoglobulin production *in vitro*, and impaired upregulation of PC genes upon stimulation. These effects were restored by inupadenant (EOS100850), a potent and highly selective small molecule A2AR antagonist. Spatial transcriptomics analysis of tumor biopsies from patients treated with inupadenant revealed that ASCs specifically increased in tertiary lymphoid structures. Altogether, these data demonstrate that A2AR plays a key role in adenosine-mediated inhibition of B cell maturation toward ASCs through a B cell-intrinsic mechanism, and that this effect is fully reverted by inupadenant.

## Introduction

Adenosine is a ubiquitous molecule that plays a crucial role in various physiological and pathological processes. It is generated in response to proinflammatory stimuli, such as cellular stress initiated by hypoxia or ischemia. Following the release to the extracellular space, adenosine triphosphate (ATP) undergoes conversion to adenosine by the two ectonucleotidases CD39 (ENTPD1) and CD73 (NT5E). Additionally, the non-canonical adenosinergic pathway metabolizes NAD+ to adenosine through the action of CD38 and CD203a/PC-1 (1).

Extracellular adenosine can activate four distinct G-protein-coupled cell-surface receptors, namely A1, A2A, A2B and A3 (encoded by ADORA1, ADORA2A, ADORA2B and ADORA3, respectively), allowing for diversity and cellular specificity in signaling pathway activity (2). The A2A and A2B receptors (A2AR and A2BR) stimulate G_s_-dependent adenylate cyclase (AC)-driven cAMP accumulation, enabling signal transduction via cAMP-activated protein kinase A (PKA). PKA catalyzes the phosphorylation of cAMP response element-binding protein (CREB) at serine 133. Once phosphorylated, phospho-CREB (pCREB) forms a dimer that binds to promoter regions of cAMP-inducible genes and allows to modulate gene expression. Activation of A2AR expressed by immune cells, including T cells and NK cells (3,4), promotes an anti-inflammatory response and reduces tissue injury in conditions of cellular stress, enabling maintenance of tissue homeostasis. The persistence of increased adenosine concentrations beyond the acute-injury phase can, however, become detrimental to tissues by activating pathways that trigger immunosuppression or promote an unremitting wound-healing process.

Landmark studies by Ohta and colleagues have highlighted the importance of adenosine for tumor immune escape (5). Within the tumor microenvironment (TME), elevated extracellular adenosine dampens the antitumor response both directly by inhibiting anti-tumor effector lymphocytes, and indirectly by affecting antigen presenting cells as well as lymphoid and myeloid regulatory cells. Until recently, studies in the field of cancer immunotherapy have focused on T cells as key players in the control of tumor development, including studies addressing adenosine as a regulatory mechanism for the inhibition of cytotoxic anti-tumor immune responses. Thus, while the effects of adenosine signaling through A2AR in T cells have been extensively characterized, less is known for other immune cells.

In the last few years, a key role of B cells and plasma cells in anti-tumor immunity has begun to emerge. The presence of tumor-infiltrating B cells, including antibody-secreting cells (ASCs), and of tertiary lymphoid structures (TLSs) is associated with better prognosis and improved response to immune checkpoint blockade in several cancers (6–8), including lung (9,10), melanoma (11), and sarcoma(12). TLSs are ectopic organized lymphoid aggregates of B cells, T cells and dendritic cells that form in the TME and progress through distinct maturation stages (13), and mature TLSs characterized by the presence of a germinal center (GC) have been identified as drivers for *in situ* B cell maturation and generation of antibody-secreting plasma blasts (PBs) and plasma cells (PCs) (14,15). Whether high extracellular adenosine in the TME affects B cell differentiation to PBs and PCs has not been investigated.

In this study, we aimed to explore the effects of adenosine on B cells. Frequencies of ASCs were reduced in adenosine-rich tumor regions, and A2AR triggering impaired B cell differentiation to ASCs and immunoglobulin production *in vitro*, an effect fully restored by the specific A2AR antagonist inupadenant. Inupadenant also boosted ASCs in TLS areas of cancer patients, suggesting a novel mechanism of action for A2AR antagonism in immunotherapy approaches.

## Materials and Methods

### Human specimens

Venous blood from healthy volunteers, all of whom signed an informed consent approved by the Ethics Committee (FOR-UIC-BV-050-01-01 ICF_HBS_HD v.5.0), was obtained by Centre Hospitalier Universitaire Tivoli, La Louviere, Belgium. Buffy coat from volunteers, all of whom signed an informed consent approved by the Ethics Committee (Contract - DOC-13116[C] / ICF English - DOC-06934[R]), was obtained by Red Cros Vlaanderen, Mechelen, Belgium.

Tonsil samples were collected from adult donors undergoing tonsillectomy and provided by DLS according to the protocol (DLS-BB044-V.1) approved by National Bioethics Committee of Medicine and Medical Devices, Romania.

Tumor biopsies for immunohistochemistry (IHC), Nanostring and Visium analysis were collected from cancer patients participating to the Phase I clinical study NCT03873883. Tumor resections for multiplex immunofluorescence, QMSI and GeoMx analysis were obtained from commercial providers (Fidelis, BioPartners, Conversant Bio and Amsbio). All samples were collected following Ethical committee approval.

### QMSI of adenosine in tumor tissue and GeoMx analysis

Tumor tissues from 10 cancer patients, stored at −80°C, were placed inside a cryostat with the temperature maintained at −20°C, and serial tissue sections were obtained for each sample. Adenosine was quantified using QMSI as previously described (33). RNA scope assay was used to verify RNA integrity on two serial sections (negative and positive control probes). For whole transcriptomic analysis (WTA), an additional slide was incubated with assay probes and stained with the morphological markers anti-PanCK, anti-CD45 and Syto13 (Nanostring). Regions of interest (ROIs) were selected based on adenosine content and segmented into areas of illumination (AOIs) based on PanCK expression. Library preparation and sequencing was performed according to the manufacturer instructions. SpatialDecon algorithm was used to estimate mixed cell type abundance in the regions of spatially-resolved gene expression studies. Within each tumor, estimated immune cell frequency was compared in adenosine-low and -high ROIs. The study was performed at Aliri (Loos, France).

### Isolation of B cells from tonsils and peripheral blood

Tonsils were sectioned into small pieces and then mechanically dissociated to a single cell suspension. Tonsillar and peripheral blood mononuclear cells (TMNCs and PBMCs, respectively) were isolated by lymphoprep density gradient centrifugation using SepMate 50 tubes (STEMCELL Technologies).

B cells were enriched from TMNCs using EasySep™ Human Pan-B Cell Enrichment Kit (STEMCELL Technologies) following the manufacturer’s protocol, and cultured as described below before single cell RNA sequencing (scRNAseq). Alternatively, TMNCs were stained for 30 min at 4°C (Supplementary Table 1), washed and resuspended at 10x10^6^ cells/ml in FACS buffer before sorting of naïve, memory, PBs and PCs B cell subsets (Supplementary Figure 3) using a FACS Aria III cell sorter (BD Biosciences). Sorted B cell subsets were analyzed using immunocytochemistry (ICC).

B cells were isolated from PBMCs using the EasySep™ Human Memory B Cell Isolation Kit (StemCell) according to the manufacturer’s instructions, and differentiated to PBs and PCs (see “In vitro differentiation of B cells” section).

### Tonsillar B cell culture and scRNAseq analysis

Tonsillar B cells were cultured for 3 h in a 24-well plate (2x10^6^ cells/ well) in X-vivo medium supplemented with 5% inactivated human serum (Biowest), 1 mM of sodium pyruvate (Na-Pyr), 1 mg/ml of CpG oligodeoxynucleotide (InvivoGen) and 5 mM CGS-2160 (Sigma) ± 300 nM inupadenant (EOS100850). Treated cells were then fixed (Fixation of Cells & Nuclei for Chromium Fixed RNA Profiling,10X Genomics Inc.) and library preparation was performed following Gene Expression Flex kit (10X Genomics Inc.) instructions. Libraries were sequenced (NovaSeq 6000) and Cellranger was used to process raw data and aligning reads to the human genome in order to generate gene counts. Data was filtered for quality, normalized, and analyzed in R using Seurat. Cells were clustered using the Louvain algorithm and visualized via UMAP, then annotated and confirmed as B-cell subtypes using reference data and marker genes. The samples were then integrated using reciprocal PCA and Seurat functions. This resulted in a unified, reprocessed dataset focusing on six major B-cell clusters. See Supplementary Methods for detailed method description.

### Immunocytochemistry of sorted B cells

Sorted B cell subsets were resuspended in PBS, and 10–30x10^3^ cells/cm^2^ were seeded for 1 h on poly-L-lysine coated slides (BosterBio #AR1065), fixed in 4% paraformaldehyde for 20 min and washed twice with PBS. A2AR ICC staining was performed using a Ventana Discovery Ultra platform (Ventana, Roche Diagnostics). Briefly, heat-induced epitope retrieval was performed in Tris-based buffer, pH 6, at 96 °C for 32 min (RiboCC, Ventana) followed by endogenous peroxidase inhibition for 8 min (DISCOVERY Inhibitor, Ventana). Cells were stained with anti-A2AR (7F6-G5-A2, Novus Biological) diluted in Dako Real antibody diluent (Agilent Technologies), detected with anti-Ms HQ/ anti-HQ HRP (Ventana) and visualizedwith DISCOVERY HRP kit. Stained samples were counterstained for 4 min (Hematoxylin II, Ventana) and digitized using Nanozoomer S60 (Hamamatsu) whole slide scanning device. The A2AR expression were calculated using a cell segmentation AI App developed using Visiopharm.

### Multiplexing immunofluorescence

Formalin-fixed paraffin embedded (FFPE) blocks from one tonsil and nine lung tumors were sectioned at 4 µm and analysed by multiplexing immunofluorescence for the expression of Pancytokeratin (PCK), A2AR, MUM-1, CD19, CD3, CD11c, CD123, and CD38 in areas of 6 mm^2^ over 39 regions of interest. Staining was performed with a Leica Bond RX instrument using OPAL™ reagents and OPAL™ tyramide immunohistochemistry (IHC) technology (Akoya). Images were captured using a Vectra Polaris (Automated Quantitative Pathology Imaging System, Akoya), and 20Xresolution (0.5 μm/pixel) images encompassing the whole slide were acquired using the DAPI, FITC, Cy3, Texas Red and Cy5 channels. Multispectral images were unmixed using inForm® software v6.4.2 (Akoya Biosciences) based on a library created from single colour controls and the autofluorescent signal was subtracted. Following unmixing, multispectral images were analyzed with HALO® image analysis platform (Indicalabs) using Highplex FL module. Lung cancer tissue was segmented in tumor and stroma areas using Random Forest-based machine learning and then the different immune phenotypes were extracted. The study was performed at Precision for Medicine (Royston, UK).

### Immunohistochemistry and Nanostring

FFPE tumor biopsies collected at baseline from 36 cancer patients participating to the Phase I clinical study NCT03873883 were sectioned and assessed for A2AR expression and gene expression using IHC and Nanostring technology, respectively. For IHC, 4 µm sections were stained with anti-A2AR antibody as described in “Immunocytochemistry” section. Stained slides were scanned and analyzed with Visiopharm software to determine the density of A2AR+cells (cells/mm^2^) in the tumor areas. For gene expression analysis, RNA was extracted from macrodissected tumor areas using High Pure FFPET RNA extraction kit and quantified using Quant-iT RiboGreen RNA Reagent and Kit. Total RNA (100 ng) was assayed using a customized nCounter PanCancer IO360 panel, and gene expression was analyzed using QCed, normalized data. The study was performed at CellCarta (Antwerpen, Belgium). Samples were grouped based on median A2AR+ cell infiltration (8.5 cells/mm^2^), and differential gene expression analysis was performed.

### CREB phosphorylation assay

Peripheralblood from healthy donors (n=4) was stimulated for 45 min at 37°C with DMSO as control, or CGS-21680 (5 μM) in the presence or in the absence of inupadenant (10 nM). After stimulation, samples were incubated at RT for 10 min with PROT1 buffer (SmartTubes) and immediately transferred at -80°C. Samples were thawed for 10 min in a cold water bath (12-14°C) and treated with Thaw/Lyse buffer according to the manufacturer’s instruction before being processed for flow cytometry. Cells were permeabilized with Phosflow Perm II buffer (BD Biosciences), stained with a solution of Brilliant Staining Buffer (BD Biosciences) and relevant antibodies (Supplementary Table S1) for 60 min, washed twice and resuspended in FACS buffer before acquisition on a LSRFortessa FACS machine (BD Biosciences).

### Cyclic AMP (cAMP) assay

The potency of inupadenant on inhibition of cAMP production was determined in HEK293-hA2AR cells (PerkinElmer) stably expressing the human A2AR. Cells were pre-incubated for 10 min with increasing concentrations of inupadenant before adding the corresponding EC90 concentration of the A2AR agonist NECA (3 nM) for 30 min, and cAMP concentration was determined using a homogeneous time-resolved fluorescence resonance energy transfer (TR-FRET) immunoassay (LANCE Ultra cAMP Kit TRF0262, PerkinElmer). The study was performed at iTeos (Gosselies, Belgium), andIC50 was calculated from 4 independent experiments, each run in duplicate.

Similarly, CHO-K1-hA1, H293-hA2B and CHO-K1-hA3 were used to determine the potency of EOS100850 on human A1, A2B and A3 receptors, respectively, using cAMP homogeneous time resolved fluorescence cellular (HTRF) functional assays. Cells were pre-incubated with increasing concentrations of inupadenant (10 nM - 30 µM) before adding the corresponding EC90 concentration of the reference agonists (20 nM CPA, 20nM NECA and 177nM IB-MECA for A1, A2B and A3 receptors, respectively). The study was performed at Euroscreen (Gosselies, Belgium), and IC50 were calculated from 3 independent experiments, each run in triplicate.

### *In vitro* differentiation of B cells

A 3-step method for in vitro differentiation of peripheral blood memory B cells was adapted from Jourdan et al (18). The medium used all along the culture was IMDM medium (Capricorn) supplemented with 10% FBS-HI (Gibco), 1% Penicillin/Streptomycin (P/S, Capricorn), 50 ug/ml Transferrin (Sigma) and 5 ug/ml Insulin (Sigma), here referred as complete IMDM medium. Cells were cultured in 24-well plates at 37 °C humidified culture incubator. On day 4, 7 and 10, treated cells were harvested, washed and analyzed by flow cytometry, while supernatants were collected and stored at -80 °C until further analysis. Untreated cells were used for next differentiation step.

Step 1 (day 0-4), expansion phase: after purification, memory B cells were seeded at 7.5 x 10^4^ to 1.5 x 10^5^ cells/well and cultured in complete IMDM medium supplemented with 20 U/ml IL-2 (Miltenyi), 50 ng/ml IL-10 (Peprotech), 10 ng/ml IL-15 (Peprotech), 10 µg/ml CpG (Invivogen) and 50 ng/ml CD40L (Miltenyi). Some wells were treated with 5 µM CGS-21680 ± 300 nM inupadenant or DMSO as negative control.

Step 2 (day 4-7), PB differentiation: on day 4, untreated cells were harvested, washed and seeded at the concentration of 2 x 10^5^ cells/well in complete IMDM medium supplemented with 20 U/ml IL-2, 50 ng/ml IL-10, 10 ng/ml IL-15 and 50 ng/ml IL-6 (Miltenyi). Some wells were treated with 5 µM CGS-21680 ± 300 nM inupadenant or DMSO as negative control.

Step 3 (day 7-10), PC differentiation: on day 7 untreated cells were harvested, washed and seeded at 2.5 x 10^5^/well in complete IMDM medium supplemented with 10 ng/mL IL-15, 50 ng/ml IL-6 and 500 UI/ml IFN-α (Miltenyi). Some wells were treated with 5 µM CGS-21680 ± 300 nM inupadenant or DMSO as negative control. and incubated for 3 days at 37 °C humidified culture incubator.

### Flow cytometry

Cells were centrifuged at 450*g* for 5 min (used for all subsequent centrifugation steps), and the supernatant was discarded. Cells were resuspended in PBS containing fixable viability dye for 20 min at 4 °C, washed in FACS buffer (PBS with 2 mM EDTA and 0.1% BSA) and incubated with human Fc block (BD Biosciences). After 10 min at room temperature, a solution of Brilliant Staining Buffer (BD Biosciences) and relevant antibodies (Supplementary Table 1) was added. After an additional 30 min of incubation at 4 °C, cells were centrifuged and resuspended in FACS buffer. Samples were analyzed on a LSRFortessa Cell Analyzer (BD Biosciences) using FACSDiva software (v.9.0.1; Becton, Dickinson and Company). Analysis of FACS data was performed using FlowJo (BD Biosciences, v.10.8.1).

### LEGENDPLEXplex analysis

Immunoglobulins were analysed in supernatants collected on day 4, day 7 and day 10 using the LEGENDplex™ Human Immunoglobulin Isotyping Panel (8-plex) kit according to the manufacturer’s instruction (Biolegend). The samples were run on a LSRFortessa (BD Biosciences) and analysed using Biolegend software (Qognit.com).

### Spatial transcriptomics (Visium) analysis

Biopsies from 5 cancer patients (head and neck n=1, melanoma n=3, lung n=1) enrolled in NCT03873883 clinical study were collected at baseline and after three weeks of inupadenant therapy. Spatial transcriptomic data were generated using the 10X Genomics Visium platform and processed with SpaceRanger. Tissue sections were pathologist-annotated as tumor or stroma. After QC, normalization, and feature selection using Seurat R package, individual Seurat objects were aligned via canonical correlation analysis. Differential gene expression and module score analyses between pre- and post-treatment samples were performed using a linear mixed-effects model to account for patient variability. See Supplementary Methods for detailed method description.

### Statistics

Statistical analysis was performed with GraphPad Prism (v.10.0.2) using one-way or two-way ANOVA with correction for multiple comparisons and matching for samples as appropriate and indicated in the relevant figure legends.

### Data Availability

Single cell and spatial transcriptomics sequencing data have been submitted to the ArrayExpress database (https://www.ebi.ac.uk/biostudies/arrayexpress), with the following identifiers: E-MTAB-15419 (*In vitro* B-cells experiment), E-MTAB-15430 (10X Visium data on tumor biopsies). Additionally, CellRanger and SpaceRanger output files and Seurat annotated objects for the 10X Visium and single cell transcriptomics datasets have been archived on Zenodo (Visium: https://doi.org/10.5281/zenodo.16365257, Single cell: https://doi.org/10.5281/zenodo.15920838).

## Results

### Plasma cell frequency is reduced in adenosine-rich tumor regions

We exploited quantitative mass spectrometry imaging (QMSI) followed by GeoMx digital spatial profiler (DSP) to evaluate the potential impact of adenosine on immune cell infiltration in human tumor tissues with spatial resolution.

We measured adenosine levels in 10 tumor tissues from six cancer indications using QMSI (Figure 1A). Within each tumor sample, we identified regions of interest (ROIs) with low and high adenosine levels to encompass intra-tumor heterogeneity (Figure 1A, bottom panels and Supplementary Figure 1A), and we performed whole transcriptomics analysis with GeoMx DSP. Spatial cell deconvolution revealed that PC frequency was significantly lower in adenosine-high ROIs (Figure 1B and 1C), especially in PanCK-stromal areas (Figure 1D). The distribution of other immune cells, including naïve and memory B cells (Figure 1C), T, NK and myeloid cells (Supplementary Figure 1B and 1C) seemed less affected by adenosine. Therefore, we investigated how adenosine directly affects PCs *in vitro* and in cancer patients.

**Figure 1:**
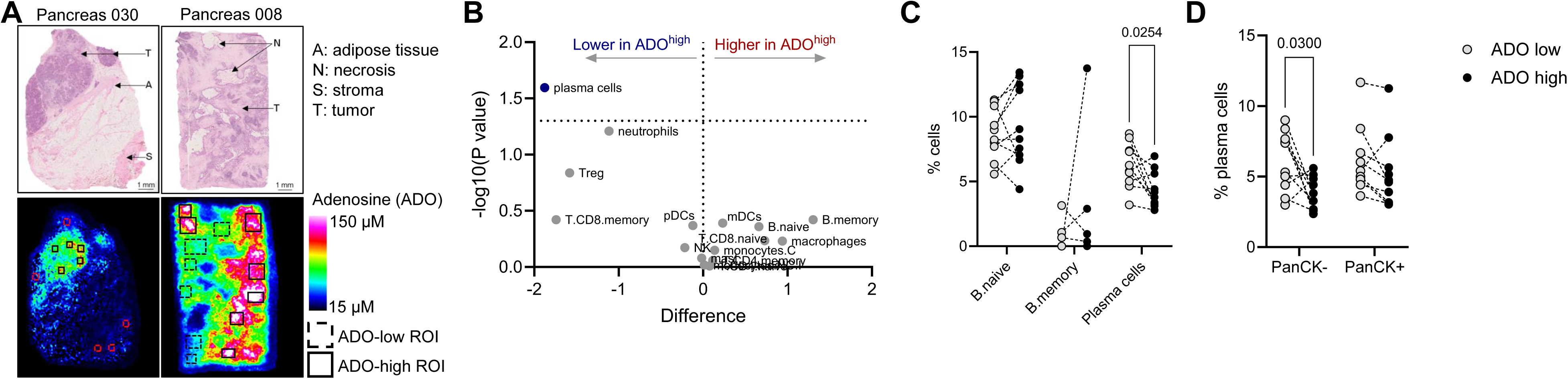
Plasma cell frequency is reduced in adenosine-rich tumor regions. Resected tumor samples (N=10) were frozen and adenosine levels determined by QMSI. GeoMx analysis was performed on selected regions of interest (ROI). **A**: Representative images of tumor sections with low (left) or high (right) adenosine levels. Top: hematoxylin and eosin staining. Bottom: QMSI images with selected ROIs for GeoMx. **B**: Volcano plot showing intra-tumor differential immune cell infiltration in adenosine-low and –high ROIs as assessed by GeoMx. **C**: Estimated percentages of B cell subpopulations in adenosine-low and –high ROIs. **D**: Estimated percentages of plasma cells in PanCK- and PanCK+ areas of adenosine-low and –high ROI. **C, D**: Each symbol represents one sample. P values from paired t test. Only significant p values (p<0.05) are displayed.

### A2A is the main adenosine receptor expressed by B cells and ASCs

To assess the expression of adenosine receptors across different B cell subpopulations, we performed single cell RNA sequencing (scRNAseq) using the 10X Genomics Chromium 3′ scRNA-seq platform on sorted and *in vitro* activated human tonsillar CD19+ B cells, and we identified six distinct B cell subsets: naïve, activated naïve, GC, cycling, memory and ASCs, the latter including short-lived PBs and long-lived PCs (Supplementary Figure 2A). Of the four adenosine receptor genes across the different B cell subpopulations, we observed almost exclusive expression of ADORA2A (Figure 2A), and among the different B cell subsets, GC B cells, ASCs and activated naïve B cells displayed highest ADORA2A expression. Immunocytochemistry (ICC) of sorted tonsil B cell subpopulations (Supplementary Figure 2B) confirmed that a significant proportion of ASCs (approximately 20% of PBs and 40% of PCs), expressed A2AR. In contrast, naïve and memory B cells exhibited minimal expression of A2AR (Figure 2B).

**Figure 2:**
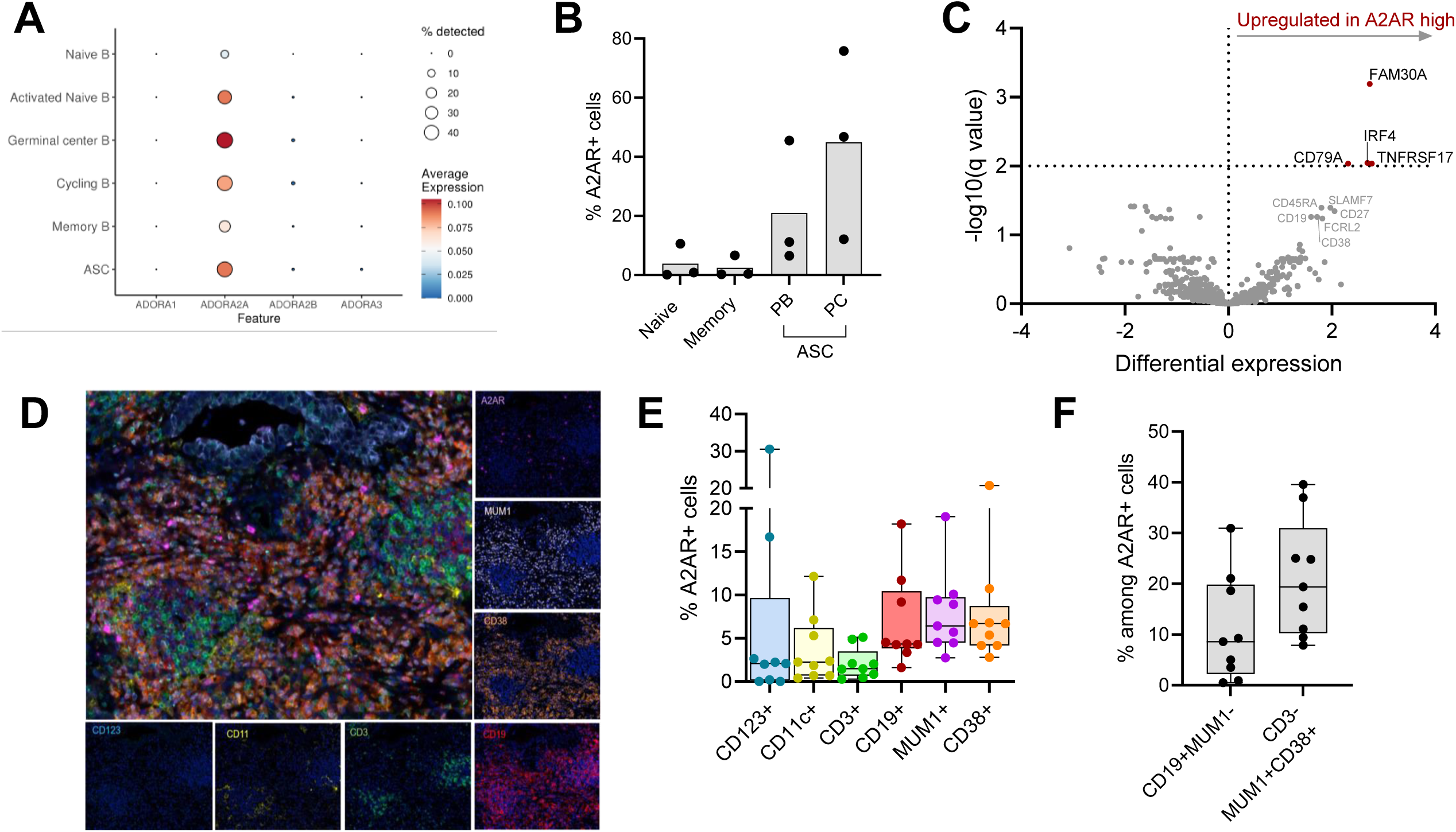
A2A is the main adenosine receptor expressed by B cells and antibody-secreting cells. **A**: Average expression of ADORA genes across different B cell subpopulations in a human tonsil by scRNAseq (DMSO condition is shown). **B**: Percentage of A2AR+ cells in sorted tonsillar B cell populations (N=3), as detected by immunocytochemistry. **C**: Volcano plot showing differentially expressed genes in tumor samples (N=36) with low or high A2AR+ cell infiltration. **D-F**: Multiplex immunofluorescence of lung cancer tissue sections (N=9). **D**: Representative images of individual and merged staining for A2AR, CD123, CD11c, CD3, CD19, MUM1, CD38. **E**: Percentage of A2AR+ cells in the indicated cell populations. **F**: Percentage of B cells (CD19+MUM1-) and bona fide antibody-secreting cells (CD3- CD19+/-MUM1+CD38+) among A2AR+ cells. **E, F**: Box plot showing median and quartiles, with whiskers from min to max. Each symbol represents one sample.

To assess if the same pattern of expression was conserved in the TME, we analyzed a public dataset of scRNAseq from melanoma tissues (11), and we confirmed that tumor-infiltrating B cells almost exclusively expressed ADORA2A, at highest levels in GC B cell-like and PC-like cells (Supplementary Figure 2C). Next, we evaluated the expression of A2AR and profiled gene expression of 36 tumor biopsies collected from metastatic cancer patients (Figure 2C). In biopsies with enriched A2AR+ cell infiltration, we observed significantly higher expression of genes involved in B cell activation, maturation and differentiation to antibody-secreting cells (FAM30A, IRF4/MUM1, TNFRSF17/BCMA and CD79A). To further identify A2AR-expressing cells in the tumor environment, we used eight-color multiplex immunostaining (A2AR, CD123, CD11c, CD3, CD19, MUM1, CD38 and PCK) (Figure 2D). As previously reported, A2AR was expressed at variable degrees in dendritic cells, T cells and B cells (16). Interestingly, a higher proportion of CD19+ B cells, MUM1+ cells and CD38+ cells expressed A2AR (Figure 2E), and B cells (CD19+MUM1-) and ASCs (CD3-cells co-expressing MUM1 and CD38) represented a substantial proportion among infiltrating A2AR+ cells (Figure 2F and Supplementary Figure 2D).

Taken together, these data indicate that A2AR is the adenosine receptor preferentially expressed by B cells and ASCs from healthy donors as well as cancer patients, prompting us to address the impact of A2AR signaling on their biology.

### Adenosine inhibits ASC differentiation and inupadenant reverts A2AR-mediated inhibition of ASCs

Although A2AR-mediated CREB phosphorylation has been reported in B cells (17), the effect of downstream signaling has not been previously well characterized.

First, we wanted to confirm that we were able to specifically trigger and inhibit A2AR signaling in our experiments. We used the prototypical A2AR agonist CGS-21680 to activate A2AR, and inupadenant (formerly EOS100850, Figure 3A), a potent and selective A2AR inhibitor (Figure 3B) to antagonize A2AR signaling. When peripheral blood was incubated with CGS-21680, we observed approximately 2-fold induction of CREB phosphorylation in CD4+ and CD8+ T cells, as well as in CD19+ B cells. Co-incubation with inupadenant (10 nM) strongly reduced CREB phosphorylation in T and B lymphocytes (Figure 3C). These results confirm that CGS-21680 triggers A2AR signaling in T and B cells, and that inupadenant potently inhibits the downstream pathway.

**Figure 3:**
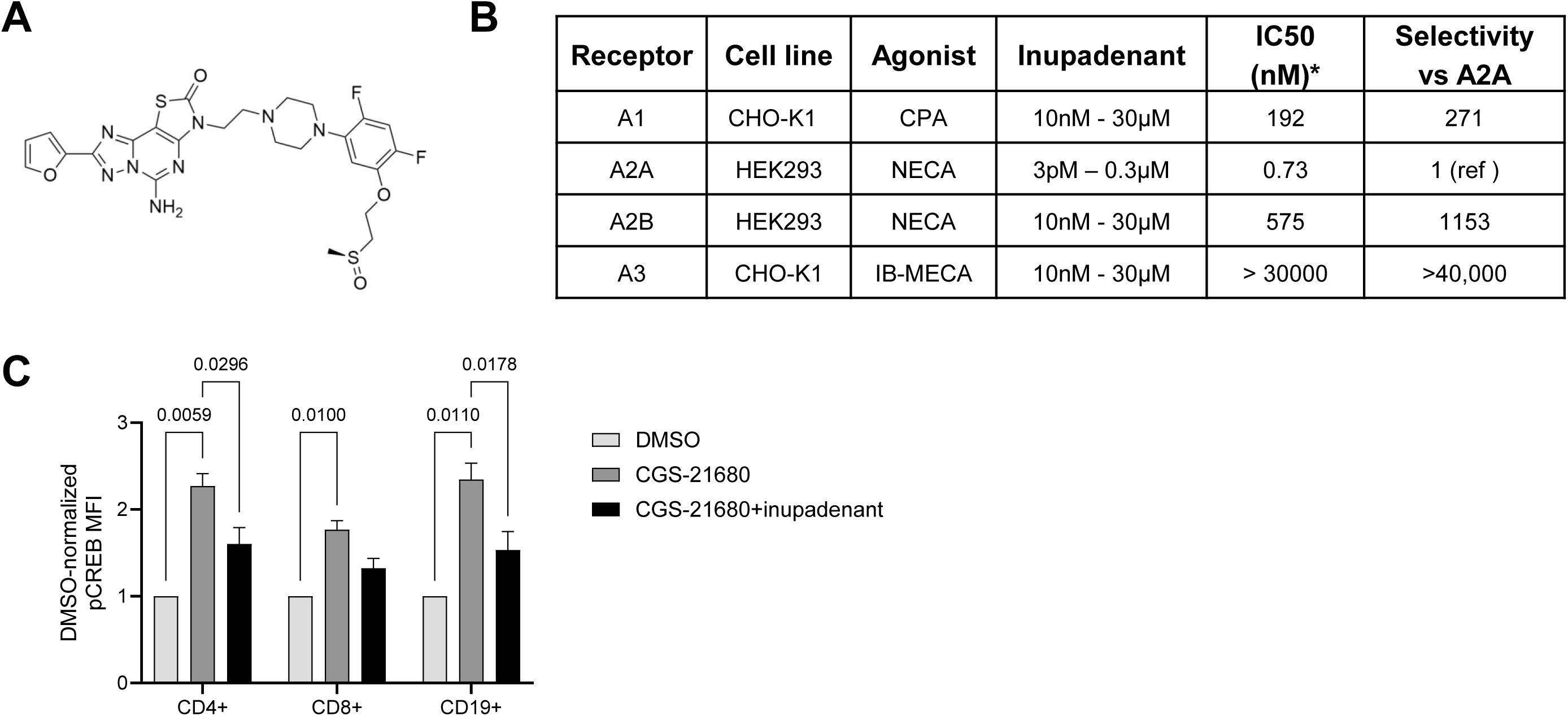
Inupadenant is a potent and selective inhibitor of A2A receptor. A: Chemical structure of inupadenant (formerly EOS100850, chemical name (+)-(S)-5-amino-3-(2-(4-(2,4-difluoro-5-(2- (methylsulfinyl)ethoxy)phenyl)piperazin-1-yl)ethyl)-8-(furan-2-yl)thiazolo[5,4-e][1,2,4]triazolo[1,5-c]pyrimidin-2(3H)-one). **B**: Potency and selectivity of inupadenant against human adenosine receptors as determined by cyclic AMP assay. **C**: Flow cytometry analysis of CREB phosphorylation in peripheral blood CD19+, CD4+ and CD8+ cells stimulated for 45 minutes with CGS-21680 (5 µM) ± inupadenant (10 nM). Data presented as mean ± SEM of N=4 healthy donors. P from ANOVA adjusted for multiple comparison. Only significant p values (p<0.05) are displayed.

PBs and PCs only represent a small fraction of B cells, limiting the characterization of modulators affecting their differentiation. Therefore, to investigate the effect of A2AR signaling in these rare populations, we used a 3-step *in vitro* assay (18) (Supplementary Figure 4A) in which CD27+ memory B cells isolated from peripheral blood were initially expanded (day 0-4), then differentiated into PBs (day 4-7) and finally into PCs (day 7-10). Flow cytometry analysis confirmed the generation of PBs and PCs after each step of the assay (Supplementary Figure 4B). CGS-21680 was added at each step of the culture in the presence or in the absence of inupadenant (Supplementary Figure 4A) and the impact of A2AR signaling on B cell viability, proliferation, maturation and differentiation into ASCs was evaluated. While CGS-21680 did not affect B cells during the expansion (data not shown), A2AR signaling significantly decreased the viability of B cells during the differentiation phases (Figure 4A), without impacting cell proliferation (Figure 4B). CGS-21680 also impaired the maturation to ASCs (Figure 4C). More specifically, the addition of CGS-21680 during the PB differentiation step (day 4-7) significantly reduced the frequency of PBs, with no effect on the small fraction of already-formed PCs (Figure 4D). Instead, when A2AR was triggered during the final differentiation step (day 7-10), the generation of PCs was affected (Figure 4E), with no effect on the PB population. Inupadenant reversed CGS-21680 effect on cell viability (Figure 4A) and completely restored A2AR-mediated suppression of B cell differentiation to ASCs, both at the level of PBs and PCs (Figure 4C-4E). These results were confirmed using an alternative method for *in vitro* differentiation of B cells into PCs (Supplementary Figure 3C-3E). Consistent with the reduced frequency of ASCs, we also observed A2AR-mediated suppression of immunoglobulin production at day 7 (Supplementary Figure 3F) and at day 10 (Supplementary Figure 3G). Once again, inupadenant inhibited the effect of CGS-21680 and restored immunoglobulin secretion.

**Figure 4:**
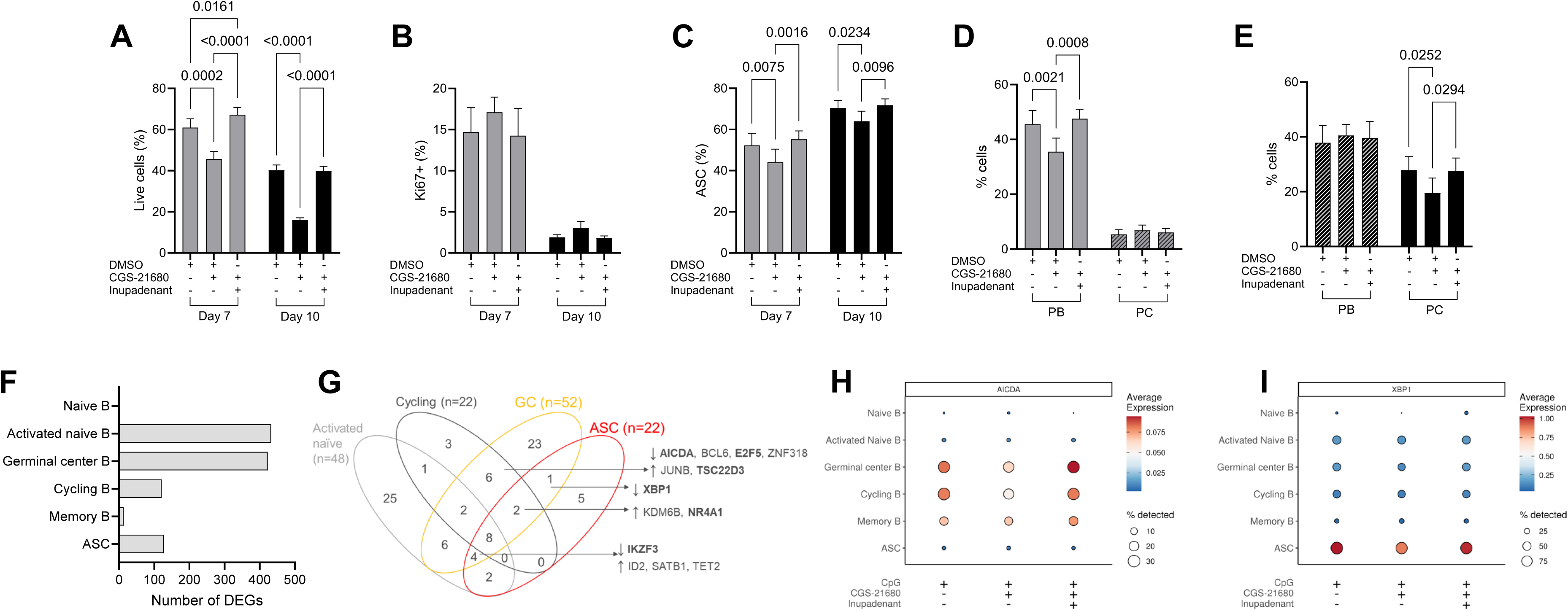
Inupadenant reverts A2AR-mediated inhibition of ASCs. A-E: CD27+ peripheral blood memory B cells were *in vitro* differentiated to plasma blasts and plasma cells, and the effects of CGS-21680 (5 µM) ± inupadenant (300 nM) were evaluated by flow cytometry. CD19+ cell viability (**A**), proliferation (**B**) and frequency of ASCs (**C**), including PBs (**D**) and PCs (**E**), at day 7 and day 10. Data presented as mean ± SEM of N=4 healthy donors. P values from one-way Anova adjusted for multiple comparison. Only significant p values (p<0.05) are displayed. **F-I**: Tonsillar B cells were stimulated for 3 hours with CpG (1 µg/ml) with or without CGS-21680 (5 µM) ± inupadenant (300 nM) and analyzed by scRNAseq. **F**: Number of differentially expressed genes in the B cell clusters observed after CGS-21860 stimulation in tonsil by scRNAseq. **G**: Venn diagram of the number of transcription factors and epigenetic enzymes modulated by CGS-21680 in the 4 most affected B cell subsets; **H, I**: Average expression of *AICDA* (**H**) and *XBP1* (**I**) across different B cell subpopulations.

These data suggest that adenosine affects B cell biology via A2AR at the level of ASC differentiation. In physiological conditions, this process occurs at the level of GC in secondary lymphoid organs and involves substantial transcriptional and epigenetic reprogramming that cannot be fully recapitulated *in vitro* (19–21). Therefore, to confirm our findings in a more physiological model, we performed an *ex vivo* assay in which CD19+ B cells sorted from human tonsils were shortly (3 hours) stimulated with CpG in the presence or in the absence of CGS-21680 and inupadenant, and scRNAseq (Supplementary Figure 2A). The highest number of differentially expressed genes (DEGs) after CGS-21680 stimulation in the six clusters of B cells (naïve, activated naïve, GC, cycling, memory and ASCs) was in the activated naïve and GC B cells (433 and 423 DEGs, respectively, Figure 4F). Cycling B and ASCs were also affected by CGS-21680 (121 and 128 DEGs, respectively), while naïve B and memory B cells were less impacted. Notably, the number of DEGs modulated by A2AR signaling in B cell clusters well correlated with their respective expression of ADORA2A (Figure 1A). This suggested that activated naïve and GC B cells were the main B cell subset affected by A2AR signaling in human tonsil. To better understand the regulatory processes activated within 3 hours of A2AR activation, we crossed our DEGs with the list of human sequence-specific DNA binding transcription factors and epigenetic regulators (TF/EE, (22)), and we identified 25 TF/EE specifically modulated in activated naïve, 23 in GC, 5 in ASCs and 3 in cycling B cells, and 32 in more than two clusters. Eight genes (HDAC9, CREM, FOSL2, NR4A2, NR4A3, STAT4, TGIF1 and ZN331) were affected in the four clusters (Figure 4G).

Among the genes modulated by A2AR signaling in the GC B cells, we observed the one for activation-induced cytidine deaminase (AIDCA), a key enzyme predominantly expressed in the GC and involved in antibody diversification through somatic hypermutation and class switch recombination (23). This gene was mainly expressed, as expected, on GC B cells (together with cycling and memory B cells). Its expression was downmodulated by A2AR signaling activation, but it was restored by inupadenant (Figure 4H). XBP1, a transcription factor essential for the development of PCs (24), was modulated in ASCs, where it was exclusively expressed (Figure 4I). Its expression was downregulated by A2AR signaling, but restored by inupadenant.

In summary, we observed that *in vitro* the transition between peripheral blood PBs and PCs and immunoglobulins production were affected by A2AR signaling. This was confirmed in the GC B cells of the tonsil, a secondary lymphoid organ that better reflects physiology. These data suggest that the activation of A2AR signaling impaired the late stage of B cell differentiation at the ASC and PC level.

### Inupadenant increases ASCs in TLS regions in patients

To elucidate the effect of inupadenant in cancer patients, we performed spatial transcriptomics analysis using Visiumplatform (10XGenomics) on paired tumor biopsies from five patients at baseline and during treatment with inupadenant across multiple cancer indications. After identification of TLS regions using the TLS signature described by Meylan et al (14), we focused on the presence of ASCs in TLS+ regions because TLSs are sites for B cell maturation and differentiation to PCs (25). Interestingly, the treatment with inupadenant increased ASCs in TLS+ regions (Figure 5A). When comparing the effect of the treatment on different immune cell signatures across regions of interest (all spots, stroma and tumor spots, total T cells and CD8+ T cell spots, B cell spots and TLS spots), the main effect of inupadenant was visible on the ASCs – in fact, the ASC signature was higher in both TLS and CD8+ T cell spots (Figure 5B), suggesting a possible interaction between these two immune cells on treatment compared to baseline.

**Figure 5:**
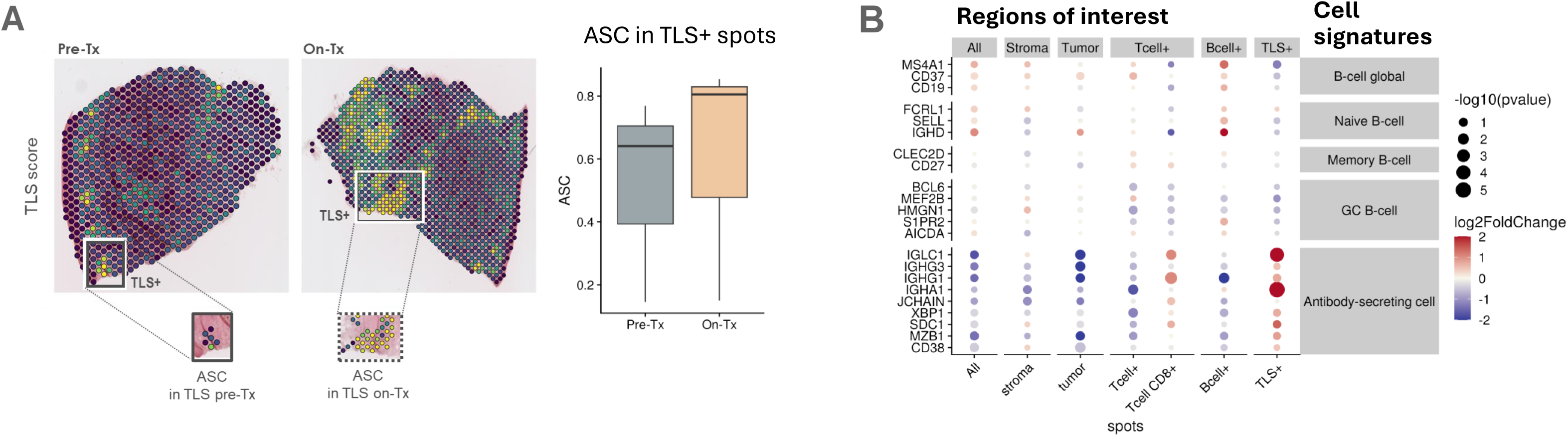
Inupadenant increases ASCs in TLS regions in patients. A: Representative tissue section showing ASCs in TLS+ regions from pre-treatment and on-treatment biopsy (left) and bar plots representing the ASCs Module Score in TLS+ spots between pre-treatment and on-treatment biopsies (right). **B**: Log2Fold change expression of different cell signatures in different regions of interest

Altogether, these data show that inupadenant treatment in cancer patients favored B cell maturation towards ASCs in the TLS regions in the tumor microenvironment.

## Discussion

Over the past decade, immunotherapy with checkpoint blockade and adoptive T cell therapy has revolutionized cancer therapy. However, although a variety of malignancies have been effectively treated with these therapies, a significant proportion of patients still fail to respond, or eventually relapse after an initial response, prompting an urgent need to identify and target other immunosuppressive mechanisms. It is now well established that the ability of the immune system to combat cancer extends beyond T cells to encompass B cells. Within the TME, elevated extracellular adenosine has been shown to suppresses immune responses by inhibiting anti-tumor effector lymphocytes, and by affecting antigen presenting cells and lymphoid and myeloid regulatory cells (2).

In this study, we present a novel mechanism for adenosine-mediated immune suppression in the TME that targets B cell maturation in an A2AR-dependent manner. After observing that the adenosine-rich regions within tumors exhibited a significantly reduced frequency of PCs, we investigated how adenosine could affect B cell biology. We found that B cells, and especially activated B cells, GC B cells and ASCs express the adenosine receptor A2AR. A2AR triggering with the selective agonist CGS-21680 inhibited B cell differentiation to ASCs *in vitro* and reduced the expression of key PC genes involved in class switch recombination and somatic hypermutation *ex vivo*. The selective A2AR antagonist inupadenant restored the expression of AICDA and XBP1 in GC/cycling B cells and ASCs, respectively, and rescued PB and PC differentiation and immunoglobulin secretion, confirming the key role of A2AR in the pathway of B cell differentiation.

Although extensively studied in other immune cell types, the role of adenosine in B cells biology has been poorly characterized. Genetic defects in adenosine deaminase (ADA) gene are associated with accumulation of adenosine in body fluids (26). Interestingly, immunophenotyping of patients with deficiencies in ADA2 (DADA2) showed defects in terminal B cell differentiation, low proportions of IgA+ and IgG+ memory cells, and impaired class switching (27). B cell intrinsic defects and defective function of T follicular helper cells contribute to DADA2 immunodeficiency (28), and a role of adenosine signaling through A2AR in dysregulation of B cell responses has been suggested (29).

In healthy subjects, it has been shown that resting peripheral blood B cells can express the adenosine-generating enzymes CD39 and CD73. In the presence of adenosine, activation and proliferation of CD39+CD73+ Bcells were affected in an autocrine manner via A3R (30). More recently, a population of CD39+CD73+ B regulatory (Breg) cells has also been described in the TME of head and neck cancer patients, while peripheral blood B effector (Beff) cells were found to predominantly express A2AR rather than A3R (31). The authors hypothesized that adenosine produced by Breg in the TME suppresses BCR signaling in Beff cells by decreasing Bruton’s tyrosine kinase phosphorylation. Our findings complement and extend this observation, and identify an additional mechanism of adenosine-mediated suppression of B cells in the TME.

Tumor infiltrating PCs are thought to be generated in the tumor TLSs. Meylan and colleagues showed that in renal cell carcinoma, mature TLSs are sites for maturation, clonal expansion, somatic hypermutation and isotype switching of B cells, providing evidence of intra-tumor differentiation of B cells (14). Similarly, in head and neck cancer, mature TLSs presented enrichment of clones from both B cells and PCs, and more expanded clonotypes of B cells, PBs and PCs, supporting the notion that TLSs are sites for anti-tumor B cell immunity in situ (15). In the present study, we show that inhibition of A2AR with inupadenant favors B cell maturation towards ASCs in the TLS regions of cancer patients. Furthermore, ASCs were also increasedin CD8+ T cell areas, suggesting a possible interaction between these two cell populations.

This study has several limitations. First, while we provide evidence for B cell intrinsic A2AR-mediated inhibition of differentiation to ASCs, we did not specifically address interactions with other cell types in this context. T follicular helper cells and dendritic cells, required to fully orchestrate the GC reaction, are also responsive to adenosine. A2AR-mediated T cell intrinsic regulation of humoral responses in GCs has been reported (32). Second, although we provide the first evidence of increased ASCs in TLS regions following A2AR blockade in cancer patients, it is important to note that this analysis included a limited number of subjects and needs further confirmation in a bigger cohort. Moreover, although a previously unappreciated role of B cells and PCs in anti-tumor immunity has begun to emerge recently, additional research is needed to determine the clinical impact of shaping humoral responses.

In summary, our study is the first to report that A2AR triggering inhibits B cell differentiation into ASCs at their terminal step. The use of inupadenant to inhibit A2AR signaling shows promise in restoring B cell functions and enhancing anti-tumor humoral responses, and deserves further testing in the clinical setting.

## Acknowledgements

We thank the patients and their families. We thank Marie-Caroline Dieu-Nosjean for her scientific support.

