## Supplementary Material for "A2AR-mediated inhibition of B cell maturation: a novel mechanism of immune suppression in cancer"

**Supplementary methods**

***In vitro* differentiation of B cells – alternative method**

Total B cells were isolated from PBMCs using B cell isolation kit II (Miltenyi Biotech) and cultured in 96-well plates for 6 days in X-VIVO15 medium supplemented with 5% human serum (Biowest), 1mM Na-Pyr (Lonza), 1x P/S (Lonza), 500 ng/ml CD40L (Human CD40-Ligand Multimer Kits, Miltenyi Biotec), 2.5 µg/ml anti-BCR (AffiniPure $F(ab')_2$ Fragment Goat Anti-Human IgA + IgG + IgM (H+L), Jackson ImmunoResearch Europe Ltd), 10 ng/ml IL-21 (R&D System) and 50 U/ml IL-2 (Miltenyi Biotec) with every 3-day passages at $4 \times 10^5$ cells/well. After 6 days, cells were cultured with or without 5 µM CGS-21680, 300 nM inupadenant or DMSO as negative control for additional 4 days before being analysed by flow cytometry.

**Single cell analysis, identification of major cell types and dataset integration**

The constructed libraries were sequenced on an Illumina NovaSeq 6000 platform with read lengths of 28 bp for Read 1 (cell barcode and UMI) and 90 bp for Read 2 (transcript). Raw FASTQ files were demultiplexed using the Cellranger mkfastq command (10X Genomics Inc.). Gene counts were generated by aligning the reads to the human reference genome (GRCh38) using the Cellranger multi pipeline. The resulting gene-barcode matrices were imported into R (v4.4.0) and analyzed using the Seurat package (v5.0.1). Quality control was performed by excluding cells with fewer than 200 features or a mitochondrial gene content over 5% and genes expressed in fewer than 3 cells. Data normalization was carried out using the *NormalizeData* function with the ‘LogNormalize’ default. The top 2000 highly variable genes were selected using *FindVariableFeatures* and used as input for the dimensional reduction through the first 30 principal components. Cells were subsequently clustered using the Louvain algorithm at a resolution of 0.5 using the *FindNeighbors* and *FindClusters* with default settings, while 2D visualization was performed using Uniform Manifold Approximation and Projection (UMAP) after PCA dimensionality reduction. For each separate sample, clusters were annotated using Azimuth (v0.5.0) based on the Tonsil v1 reference (Massoni-Badosa et al, 2022). Based on this annotation, only B-cell subtypes were retained, resulting in 6 major clusters. Identification of these subtypes was confirmed using canonical marker genes (Supplementary Table 2).

Samples were integrated into one object using reciprocal PCA (RPCA). 2000 integration anchors using the *FindIntegrationAnchors* and *IntegrateData* Seurat functions were used. The integrated Seurat object was rescaled and reprocessed with the same functions as described above.

**RNA Quality Assessment and sequencing for Visium analysis**

Biopsies from 5 cancer patients (head and neck n=1, melanoma n=3, lung n=1) enrolled in NCT03873883 clinical study were collected at baseline and after three weeks of inupadenant therapy. Up to five FFPE sections per tissue sample were used for HE staining and RNA extraction. For each sample, a capture area of 6.5 x 6.5 mm (5.000 spots) was annotated by a pathologist. RNA extraction was performed with the High Pure FFPET RNA extraction Kit (Roche). DV200 values were evaluated using BioAnalyzer (Agilent Technologies), and only samples with DV200 values higher than 30% were included in the analysis. Generated libraries were then sequenced on the Illumina NovaSeq 6000 platform. The study was performed at CellCarta (Antwerpen, Belgium).

**Spatial Transcriptomics analysis**

Spatial transcriptomic data were obtained using the 10X Genomics Visium platform and processed via Space Ranger software (v2.1.1, 10X Genomics Inc.). Raw sequencing reads were aligned to the human reference transcriptome (GRCh38; refdata-gex-GRCh38-2020-A) utilizing the 10X spaceranger count pipeline. Histological H&E-stained tissue sections corresponding to each Visium slide underwent evaluation and annotation by an experienced pathologist. Regions of interest were manually outlined and classified as either tumor or stroma based on established morphological criteria. These annotations enabled spatial contextualization of transcriptomic results and facilitated the correlation of gene expression patterns with tissue architecture.

Subsequent analyses employed the Seurat R package (v5.1.0). Quality filtering of spatial spots was conducted with the following thresholds: nCount_Spatial > 5,000, nFeature_Spatial > 500, and percent.mt < 20%, to retain high-quality data. Data normalization was performed using Seurat's NormalizeData function under default parameters. Variable features were identified with FindVariableFeatures, applying the "vst" selection method and retaining the top 2,000 genes for downstream analysis. Scaling of the data was completed using Seurat's ScaleData function.

To integrate datasets across multiple samples and experimental conditions, individual Seurat objects were aligned through canonical correlation analysis (CCA). Integration anchors were identified using FindIntegrationAnchors (with options reduction = "cca", dims = 1:30), and data sets were integrated using IntegrateData with k.weight set to 50 to balance both sample-specific and shared variation. The resulting integrated dataset was utilized for all subsequent analyses.

Gene signature scores for predefined gene sets were calculated per spatial spot using Seurat's AddModuleScore function. To evaluate treatment-related changes in gene expression and module scores, we conducted differential analyses between pre- and post-treatment samples. A linear mixed-effects model was applied to account for intra-patient variability, specified as expressionscore ~ time (pre vs post) + (1|patient_id), where 'expression' denotes gene expression, 'score' refers to the module score for each spot, 'time' reflects the treatment condition, and 'patient_id' was modeled as a random intercept to account for repeated measures within individuals. This modeling approach was implemented using the lmerTest package in R.

| Antigen | Fluorochrome | Clone | Dilution | Provider |
|---|---|---|---|---|
| **Sorting of B cell subsets** | | | | |
| CD45 | FITC | REA747 | 1:200 | Miltenyi Biotech |
| CD3 | PerCP | REAL104 | 1:100 | Miltenyi Biotech |
| CD38 | Vio Bright V423 | REA572 | 1:100 | Miltenyi Biotech |
| CD19 | APC | REA675 | 1:200 | Miltenyi Biotech |
| IgD | Vio Bright R720 | REA740 | 1:100 | Miltenyi Biotech |
| CD27 | PE | REA499 | 1:100 | Miltenyi Biotech |
| HLADR | PE-vio 770 | REA805 | 1:100 | Miltenyi Biotech |
| Viability dye | eF780 | | 1:500 | Invitrogen |
| ***In vitro* differentiation of B cells** | | | | |
| CD38 | BV421 | HIT2 | 1:100 | BD Biosciences™ |
| IgD | BV510 | IA6-2 | 1:100 | BD Biosciences™ |
| CD3 | PerCP-Cy5.5 | UCHT1 | 1:50 | BD Biosciences™ |
| CD27 | PE | L128 | 1:25 | BD Biosciences™ |
| HLA-DR | PE-Cy7 | L243 | 1:200 | BioLegend |
| CD19 | AF647 | HIB19 | 1:100 | BioLegend |
| CD19 | BV711 | HIB19 | 1:100 | BioLegend |
| Viability dye | eF780 | | 1:500 | Invitrogen |
| Ki67 | AF488 | Ki-67 | 1:50 | BioLegend |
| CD45 | BUV395 | HI30 | 1:500 | BD Biosciences™ |
| **CREB phosphorylation assay** | | | | |
| CREB/AFT-1 | PE | pS133/S63 | 1:6.25 | BD Biosciences™ |
| CD4 | BV786 | SK3 | 1:100 | BD Biosciences™ |
| CD8a | Alexa Fluor 488 | HIT8a | 1:100 | Biolegend |
| CD19 | BV711 | SJ25C1 | 1:25 | Biolegend |

**Supplementary Table 1. List of antibodies used in flow cytometry for the indicated analysis**

| B cell subtypes | Marker Genes | Reference |
|---|---|---|
| Activated naive | CD69, CCR7, CD83 | (33,34) |
| Naïve | FCMR, HVCN1, SELL, IGHD | (34) |
| Germinal center | LM02, FGD6, MME, AICDA | (33) |
| Cycling | PCNA, TUBA1B, TOP2A | (33) |
| Memory | MS4A1, KLF2 | (33,34) |
| Antibody secreting cells | XBP1, JCHAIN, MZB1, PRDM1 | (34) |

**Supplementary Table 2. List of marker genes to identify B cell subpopulations**

| Antibody | Type | Clone | Provider | Fluorophore |
|---|---|---|---|---|
| CD3 | Rabbit IgG | D7A6E | Cell Signaling Technology | OPAL520 |
| PCK | Mix of mouse IgG1 and IgG2a | - | Sigma | OPAL480 |
| CD19 | Rabbit IgG | D4V4B | Cell Signaling Technology | OPAL650 |
| CD11c | Rabbit IgG | D3V1E | Cell Signaling Technology | OPAL690 |
| CD123 | Rabbit IgG (polyclonal) | - | Thermo Fisher | OPAL540 |
| CD38 | Rabbit IgG | SP149 | Roche | OPAL620 |
| A2AR | Mouse IgG2a | 7F6-G5-A2 | Novus | OPAL570 |
| MUM-1 | Mouse IgG1 | MUM1p | Dako | OPAL780 |

**Supplementary Table 3. List of the antibodies and fluorophores used for the multiplex panel.**

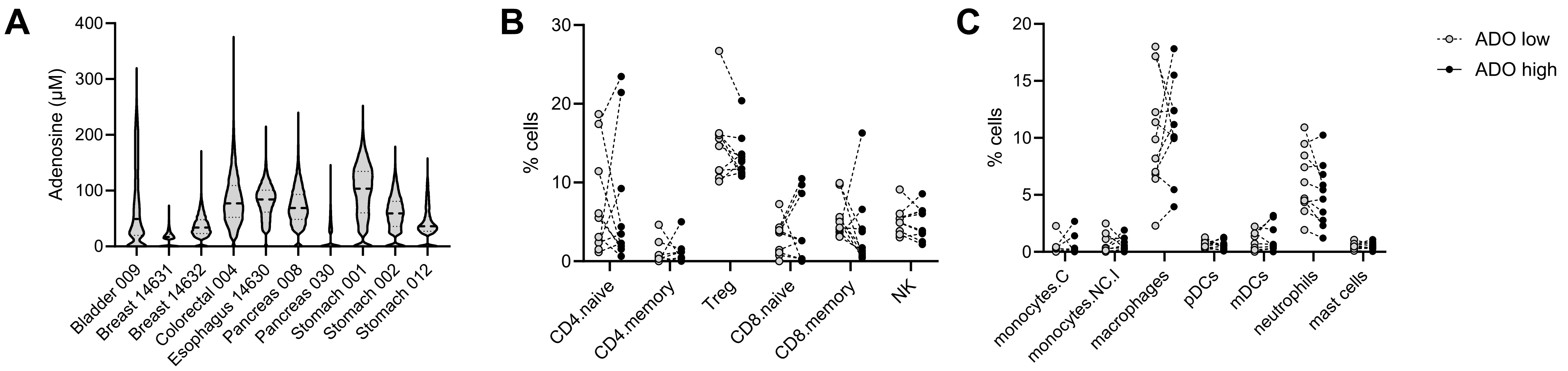


**Supplementary Figure 1. A**: Violin plot of adenosine distribution in the selected ROIs. **B, C**: Estimated percentages of lymphoid (**B**) and myeloid (**C**) cell populations in adenosine-low and –high ROIs. Each symbol represents one sample, analysis by paired t test. All p>0.05 (not displayed).

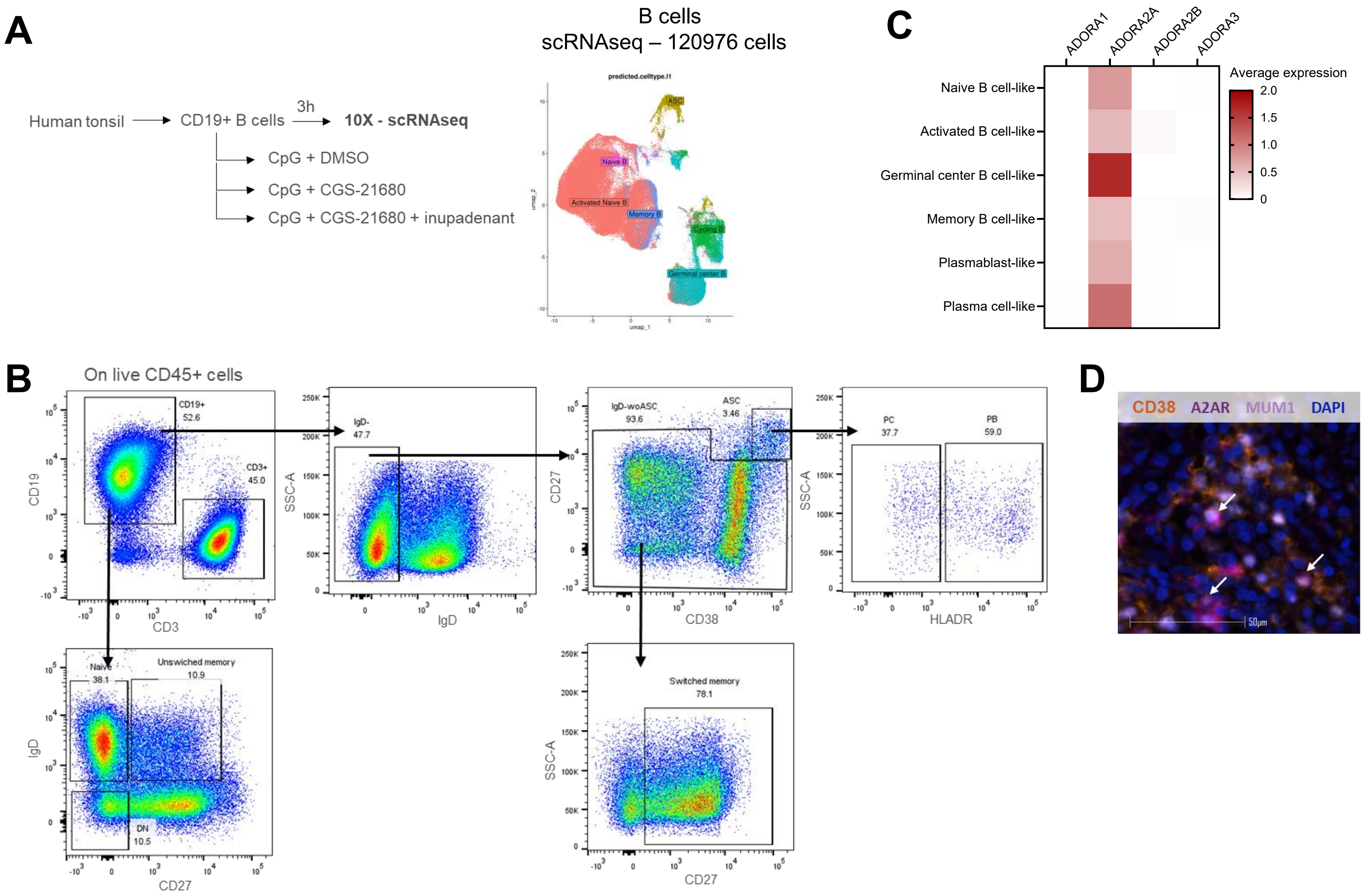


**Supplementary Figure 2. A**: Schematic representation of scRNAseq experiment (left); UMAP plot of human tonsillar B cells scRNAseq data. **B**: Gating strategy for flow cytometry analysis of different B cell populations. One representative tonsil sample is shown. ASCs (antibody secreting cells): CD19+IgD-CD38++CD27++; PB (plasma blasts): CD19+IgD-CD38++CD27++HLADR+; PC (plasma cells): CD19+IgD-CD38++CD27++HLADR-; naïve B cells: CD19+CD27-IgD+; DN (double negative): CD19+CD27-IgD-; switched memory: CD19+IgD-CD27+w/oCD27++CD38++. **C**: Average expression of ADORA genes across different B cell subpopulations in human melanomas by scRNAseq. **D**: Representative image of merged staining for A2AR, MUM1 and CD38. Arrows indicate cells positive for the three markers.

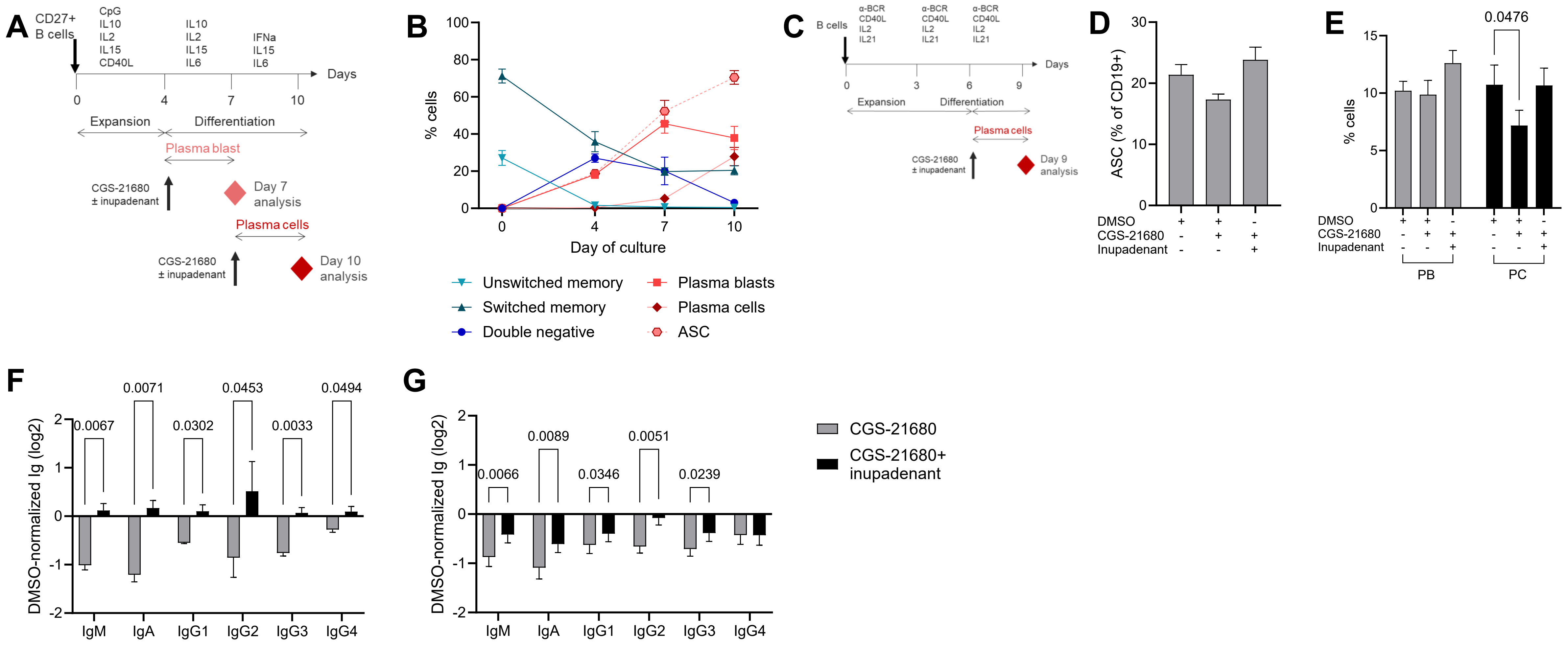


**Supplementary Figure 3. A:** Schematic representation of *in vitro* differentiation methods of peripheral blood CD27+ purified memory. **B**: Flow cytometry analysis of cultured cells in the absence of CGS-21680 and inupadenant at day 0, 4, 7 and 10. **C-E**: Schematic representation of an alternative method to differentiate peripheral blood CD19+ B cells into PBs and PCs (**C**), and flow cytometry analysis of the effect of CGS-21680 ± inupadenant on the frequency of ASC (**D**) including PB and PC (**E**) at day 10. **F, G**: Effect of CGS-21680 ± inupadenant on immunoglobulin (Ig) secretion at day 7 (**F**) and day 10 (**G**) of method described in A. Ig were quantified with Legendplex, normalized to DMSO condition and log2-transformed. Data presented as mean ± SEM of N=4 healthy donors. P from ANOVA adjusted for multiple comparison. Only significant p values ($p<0.05$) are displayed.
